# From Cohorts to Molecules: Adverse Impacts of Endocrine Disrupting Mixtures

**DOI:** 10.1101/206664

**Authors:** Lina Birgersson, Gábor Borbély, Nicolò Caporale, Pierre-Luc Germain, Michelle Leemans, Filip Rendel, Giuseppe Alessandro D’Agostino, Raul Bardini Bressan, Francesca Cavallo, Nadav Even Chorev, Vesna Munic Kos, Maddalena Lazzarin, Steven M. Pollard, Birgitta Sundström, Alejandro Lopez Tobon, Sebastiano Trattaro, Matteo Zanella, Åke Bergman, Pauliina Damdimopoulou, Maria Jönsson, Wieland Kiess, Efthymia Kitraki, Hannu Kiviranta, Mattias Öberg, Panu Rantakkoko, Christina Rudén, Olle Söder, Carl-Gustaf Bornehag, Barbara Demeneix, Jean-Baptiste Fini, Chris Gennings, Eewa Nånberg, Joëlle Rüegg, Joachim Sturve, Giuseppe Testa

**Affiliations:** Department of Biological and Environmental Sciences. University of Gothenburg, Gothenburg, Sweden.; Unit for Toxicology Sciences, Swetox, Karolinska Institutet, Södertälje, Sweden.; Department of Experimental Oncology, Laboratory of Stem Cell Epigenetics, European Institute of Oncology, Milan, Italy; Department of Oncology and Hemato-Oncology, University of Milan, Milan, Italy; UMR 7221, Evolution of Endocrine Regulations. CNRS/ Muséum National d’Histoire Naturelle, Sorbonnes Universités, Paris, France; Faculty of Health, Science and Technology, Department of Health Sciences, Karlstad University, SE-651 88 Karlstad, Sweden; Centre for Regenerative Medicine and Edinburgh Cancer Research Centre, University of Edinburgh, Scotland; Department of Organismal Biology, Uppsala University, Uppsala, Sweden; Hospital for Children and Adolescents. Department of Women and Child Health. University Hospital. University of Leipzig, Germany; School of Health Sciences, National and Kapodistrian University of Athens, Athens, Greece; National Institute for Health and Welfare. Kuopio, Finland; Department of Applied Environmental Science, Stockholm University, Stockholm, Sweden; Icahn School of Medicine at Mount Sinai, New York, USA.; Department of Clinical Neuroscience, Karolinska Institutet, Stockholm, Sweden.

## Abstract

Convergent evidence associates endocrine disrupting chemicals (EDCs) with major, increasingly-prevalent human disorders. Regulation requires elucidation of EDC-triggered molecular events causally linked to adverse health outcomes, but two factors limit their identification. First, experiments frequently use individual chemicals, whereas real life entails simultaneous exposure to multiple EDCs. Second, population-based and experimental studies are seldom integrated. This drawback was exacerbated until recently by lack of physiopathologically meaningful human experimental systems that link epidemiological data with results from model organisms.

We developed a novel approach, integrating epidemiological with experimental evidence. Starting from 1,874 mother-child pairs we identified mixtures of chemicals, measured during early pregnancy, associated with language delay or low-birth weight in offspring. These mixtures were then tested on multiple complementary *in vitro* and *in vivo* models. We demonstrate that each EDC mixture, at levels found in pregnant women, disrupts hormone-regulated and disease-relevant gene regulatory networks at both the cellular and organismal scale.

## Introduction

Human populations are exposed to a large number of chemicals with endocrine disrupting properties (EDCs)^1^. Their regulation represents a major unmet challenge due to the fact that while exposure to single EDCs has repeatedly been associated with major disorders and impaired development^2^, real life entails simultaneous exposure to multiple EDCs in mixtures, with additive effects at lower doses than experimental effect thresholds for single compounds^3^. Experimental evidence with mixtures is however most often limited to combinations within the same chemical class or to observational measurements on more complex mixtures, thus lacking causative weight to link actual population-based exposure with adverse health outcomes in humans. Here we pursued a systematic integration of epidemiological and experimental evidence to elucidate the molecular pathways affected by EDC mixtures that are causally related to adverse outcomes in humans.

## Results

### An integrated epidemiological-experimental design assessing the impact of EDC mixtures on human health and development

To assess health outcomes of real-life EDC exposures we harnessed: i) the power of a population-based mother-child pregnancy cohort to measure prenatal EDC exposures, combined with novel biostatistical tools to infer associations between specific EDC mixtures and two child health domains: neurodevelopment and metabolism/growth (Figure 1a); ii) complementary assays in human systems, to establish causality and deconvolute gene regulatory networks and *in vitro* cellular responses dysregulated by these EDC mixtures in concentrations corresponding to human exposure (Figure 1b); iii) paradigmatic *in vivo* models to determine the physiological impact of key affected pathways (Figure 1c).

**Figure 1:**
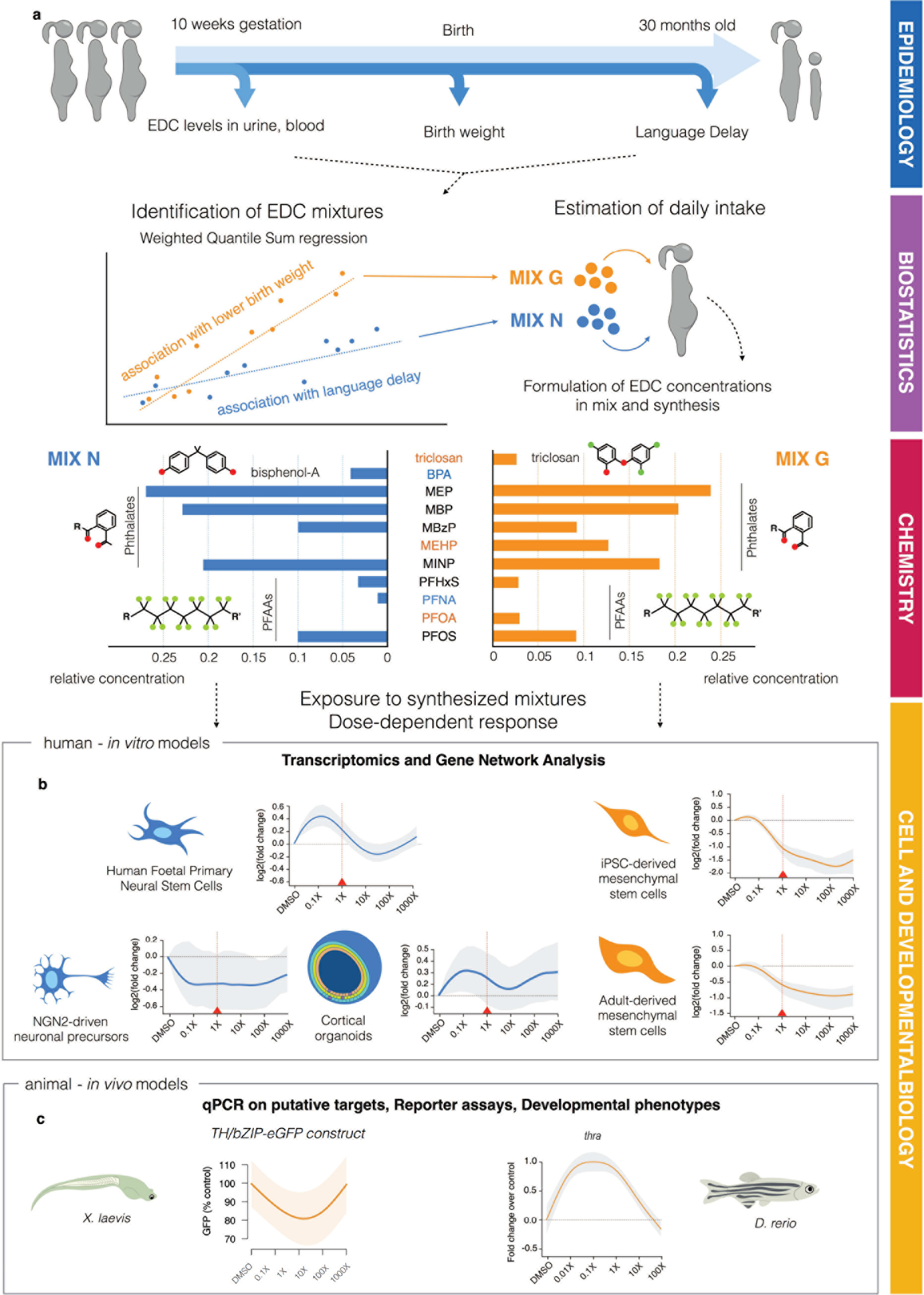
Overview of the study. **a: Identification of two EDC mixtures that are associated with adverse health outcomes in two health domains, neurodevelopment and growth**. In the SELMA pregnancy study, 20 EDCs and metabolites were measured in urine or serum of women around pregnancy week 10. Associations between these exposures and language delay of the children at age 2.5 years or birth weight were established using weighted quantile sum regression. This resulted in the identification of so-called “bad actors” that contributed to the association with the adverse health outcome (language delay or reduced birth weight) in the mixture. Based on their ratios found in the SELMA women’s serum, the identified bad actors were mixed to compose MIX N (based on the association with language delay, blue coloured) and MIX G (based on the association with birth weight, orange coloured) for subsequent use in the experimental systems in concentrations corresponding to 0.01X, 0.1X, 1X, 10X, 100X, and 1000X serum concentrations in the SELMA mothers. **b: Identification of gene regulatory networks and cellular responses dysregulated by MIX N and MIX G, along with their dose-response relationships**. Transcriptome analyses were carried out in Human Foetal Primary Neural Stem Cells (HFPNSC), NGN2-driven neural precursors, or cortical organoids as well as in iPSC-derived or adult mesenchymal stem cells (MSCs) upon 48 h treatment with 0.1-1000X MIX N and MIX G, respectively. Predominant dose-response patterns showed, in most models, non-monotonic shapes. Significant transcriptional changes were detected already at 1X concentrations. **c: Validation of key pathways affected by MIX N and MIX G and their physiological impact in paradigmatic *in vivo* models**. Effects on key genes and pathways identified in the cellular models were assessed in *Xenopus laevis* and *Danio rerio* larvae upon short-term treatment (48-72 h), exemplified by GFP levels in transgenic Xenopus bearing a thyroid hormone-induced GFP reporter (left panel) and thyroid hormone receptor alpha (*thra*) expression quantified using RT-qPCR in zebrafish (right panel) upon MIX G exposure.

### Definition and establishment of EDC-mixtures impacting human neurodevelopment and metabolism

Humans are exposed to several classes of EDCs including phthalates, phenols and perfluorinated alkyl acids (PFAAs) ^1,2^. We focused on prenatal exposures to mixtures of 15 endocrine disruptive parent compounds (comprising 20 analytes/metabolites known or suspected of being endocrine disruptors^2^ from these classes, to determine their potential associations with two major child health outcomes: neurodevelopment, measured as language delay at 30 months of age which is an early marker of cognitive development^4^, and metabolism/growth, measured as birth weight where low birth weight is associated with metabolic syndrome including obesity and glucose intolerance^5^. First we established exposure using biomonitoring data from the Swedish Environmental Longitudinal, Mother and child, Asthma and allergy (SELMA)^6^ pregnancy cohort that includes 1,874 pregnant women (Table 1a), assessed for their urinary or serum EDC levels around the 10^th^ week of gestation (Table 1b). Specifically, we profiled urine levels of 10 metabolites of 5 phthalates, bisphenol A (BPA) and triclosan (TCS), as well as serum levels of 8 perfluorinated alkyl acids (PFAAs).

**Table 1a:**
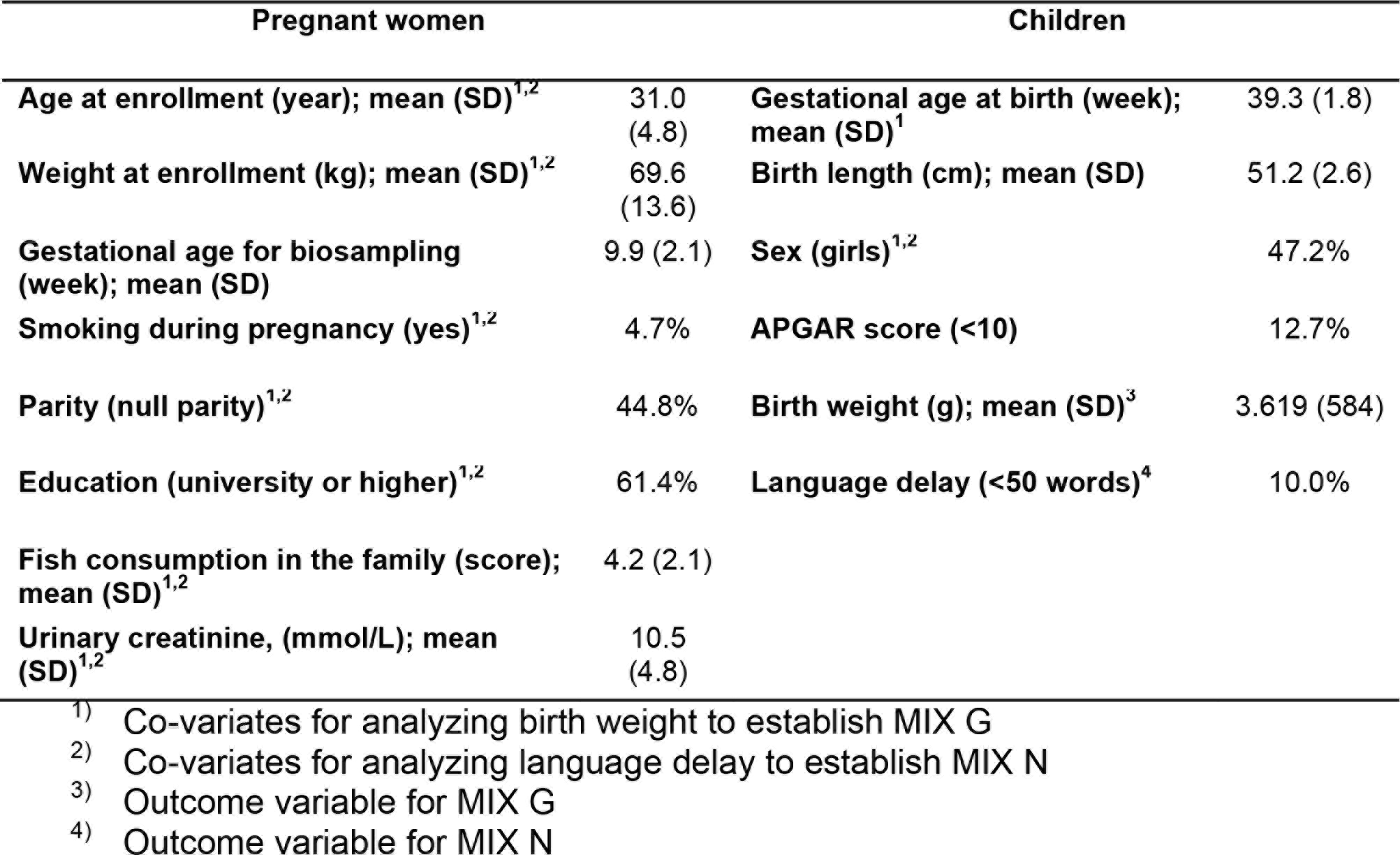
Description of the study population including 1,874 pregnant women and their children in the SELMA study.

| Pregnant women |  | Children |  |
| --- | --- | --- | --- |
| **Age at enrollment (year); mean (SD)<sup>1,2</sup>** | 31.0 (4.8) | **Gestational age at birth (week); mean (SD)<sup>1</sup>** | 39.3 (1.8) |
| **Weight at enrollment (kg); mean (SD)<sup>1,2</sup>** | 69.6 (13.6) | **Birth length (cm); mean (SD)** | 51.2 (2.6) |
| **Gestational age for biosampling (week); mean (SD)** | 9.9 (2.1) | **Sex (girls)<sup>1,2</sup>** | 47.2% |
| **Smoking during pregnancy (yes)<sup>1,2</sup>** | 4.7% | **APGAR score (<10)** | 12.7% |
| **Parity (null parity)<sup>1,2</sup>** | 44.8% | **Birth weight (g); mean (SD)<sup>3</sup>** | 3.619 (584) |
| **Education (university or higher)<sup>1,2</sup>** | 61.4% | **Language delay (<50 words)<sup>4</sup>** | 10.0% |
| **Fish consumption in the family (score); mean (SD)<sup>1,2</sup>** | 4.2 (2.1) |  |  |
| **Urinary creatinine, (mmol/L); mean (SD)<sup>1,2</sup>** | 10.5 (4.8) |  |  |

<sup>1)</sup> Co-variates for analyzing birth weight to establish MIX G
<sup>2)</sup> Co-variates for analyzing language delay to establish MIX N
<sup>3)</sup> Outcome variable for MIX G
<sup>4)</sup> Outcome variable for MIX N

**Table 1b:**
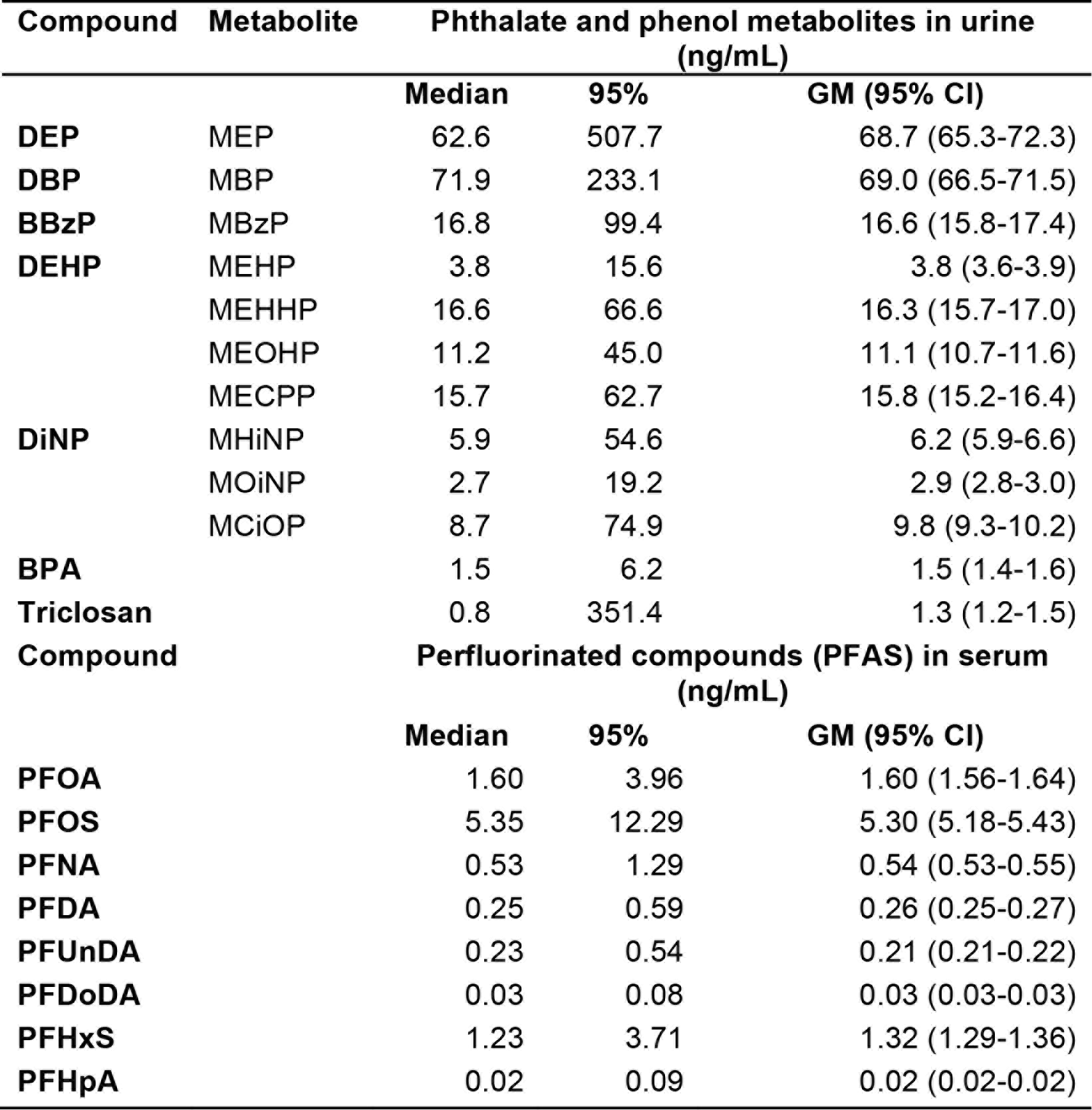
Distribution of phthalate and phenol metabolites in urine and perfluorinated compounds (PFAS) in serum analysed in 1^st^ trimester of 1,874 pregnant women in the SELMA study.

Specific mixtures (in terms of both composition and dose) were associated with the two health outcomes in a three-step procedure (Figure 1a). First, we identified the prenatal exposure to EDCs, hereafter “bad actors”, that was associated with lower birth weight or language delay in children by using weighted quantile sum (WQS) regression^7^. Next, we estimated the equivalent daily intake (DI) of “bad actors” measured in the urine (i.e., phthalates and alkyl phenols), and estimated serum concentrations from the DI for these urinary measurement-based compounds. Finally, we used the geometric means, on a molar basis, for either the measured or estimated serum levels of all compounds and established mixing proportions to prepare, for experimental validation, the two mixtures associated to language delay (MIX N) and lower birth weight (MIX G) (Figure 1a; Extended data Table 1). Mixtures were tested across concentrations (0.01X, 0.1X, 1X, 10X, 100X, 1000X) corresponding to human exposure (Table 1b), where X denotes the geometric mean of exposure levels in SELMA pregnant women.

### MIX N disrupts human neurodevelopmental pathways

To define the molecular impact of MIX N, we employed primary neural stem cells sourced from cortex and ganglionic eminence of human foetuses at post-conception week (PCW) 11 and 8, respectively (Figure 2a) (henceforth Human Foetal Primary Neural Stem Cells (HFPNSC)). Given the potentially non-linear and non-monotonic dose-response patterns associated with EDC mixtures^8,9^, the experimental design included 5 doses of MIX N, ranging from 0.1X to 1000X and a global assessment of impact on gene expression. To this end, RNA-seq was performed after 48h MIX N exposure and patterns of EDC dose-dependent transcriptional responses determined using an analysis that considers MIX N dilutions (including the DMSO control) as distinct categories. This unbiased approach, which does not assume any particular response pattern (e.g., linearity or monotony), allowed us to define lists of differentially-expressed genes (DEGs), henceforth “unbiased DEGs”, which were subsequently clustered on the basis of their dose-response patterns (Figure 2b). Next, we used the dose-response patterns to re-interrogate the transcriptomes by regressing each gene to each pattern, permitting identification of larger sets of high confidence DEGs following each pattern (Extended data Figure 1a). The dose-response patterns showed dysregulation already at low concentrations, highlighting the significance of doses recapitulating human exposure.

**Figure 2:**
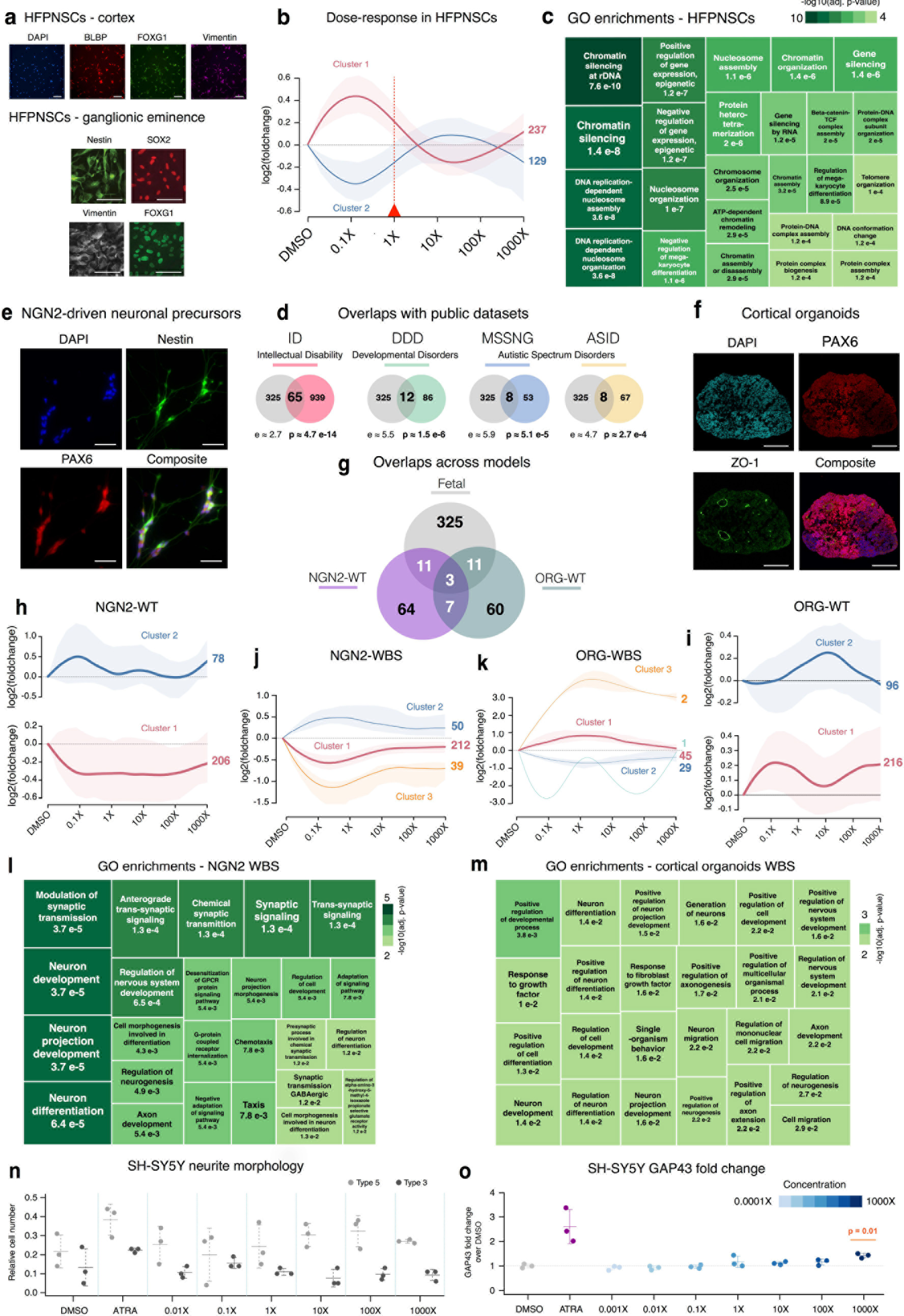
MIX N disrupts neurodevelopmental pathways in human cellular models. **a:** Immunofluorescence of paradigmatic neural stem cell markers are shown for cortical (scale bar=10μm) and ganglionic eminence (scale bar=20μm) derived human foetal primary neural stem cells (HFPNSC). **b:** Unbiased DEGs were identified through categorical analysis and were clustered into major dose-responses patterns, plotted across the different MIX N dilutions used for exposure. **c:** Treemap of the enriched gene ontology terms for the unbiased DEGs of HFPNSC (the size and color of the boxes is proportional to the significance of the enrichment). **d:** Overlaps, enrichment and significance between unbiased HFPNSC DEGs and genes associated to intellectual disability, developmental disorders and autism spectrum disorders in published databases. e: Immunofluorescence of paradigmatic neuronal markers are shown for NGN2-driven neural precursors (scale bar=20μm). **f:** Immunofluorescence of paradigmatic neuronal markers are shown for cortical organoids (scale bar=200μm). **g:** Overlaps between unbiased DEGs of HFPNSC and unbiased DEGs of the WT cell line differentiated into NGN2-driven neural precursors and cortical organoids. **h:** Unbiased DEGs, previously identified for HFPNSC, plotted in the WT cell line differentiated into NGN2-driven neural precursors across the different MIX N dilutions. i: Unbiased DEGs, previously identified for HFPNSC, plotted in the WT cell line differentiated into cortical organoid across the different MIX N dilutions. **j:** Unbiased DEGs identified through categorical analysis in the 2 WBS cell lines, differentiated into NGN2-driven neural precursors, clustered into major dose-responses patterns and plotted across the different MIX N dilutions used for exposure. **k:** For the 2 WBS cell lines, differentiated into cortical organoids, unbiased DEGs were identified through categorical analysis, clustered into major dose-responses patterns and plotted across the different MIX N dilutions used for exposure. **l:** treemap of the enriched Gene Ontology (GO) categories for the unbiased DEGs of WBS NGN2-driven neural precursors is shown. **m:** Treemap of the enriched gene ontology categories for the unbiased DEGs of WBS cortical organoids is shown. **n:** Relative number of cells with immature neurites (Type 3) or maturing neurites with nodes (Type 5) in SH-SY5Y cells treated with MIX N for 96 hours. Also shown are vehicle (0.01 % DMSO) and positive control (ATRA). Data are shown as mean ± SD of the relative amount of type 5 and 3 cells with three experimental replicates. Testing for dose-response was significant (p=0.017) in a one way ANOVA with a cubic fit on the ratio of type 5 and 3 cells. **o:** Effect of MIX N on *GAP43* mRNA expression in SH-SY5Y cells. Treatment with 10 μM ATRA was included as a positive control. Data were quantified by the relative standard curve method and expressed as the relative amount of *GAP43* divided by the relative amount of *GAPDH*. Data are shown as mean ± SD of 3 experimental replicates. Testing for dose-response was significant (p=0.001) in a one way ANOVA. The p-value in the figure is from a TUKEY HSD post hoc test on the relative amount of *GAP43* divided by the relative amount of *GAPDH* compared to vehicle (DMSO). For dose-response plots, the size of the line is proportional to the number of genes in the cluster, and shading represents 80% of the log2-fold change variation across genes.

Functional characterization of DEGs revealed enrichment in Gene Ontology (GO) categories related to chromatin modulation and regulation of gene expression (Figure 2c), showing a major and specific impact of EDC during early forebrain development. Given our epidemiological evidence associating MIX N exposure to language delay, and the central role of chromatin dysfunction in autism and intellectual disabilities^10^ we tested whether EDC-induced DEGs were enriched for genes associated with these conditions. We found significant enrichment for genes associated with Intellectual Disabilities (p ≃ 4.7 e-14 ID^11^, Developmental Disorders (p ≃ 1.5 e-6 DDD^12^) and Autism Spectrum Disorders (ASD) (p ≃ 5.1 e-5 in the Autism Speaks-Google MSSNG database^13^ and p ≃ 2.7 e-4 in the Autism Spectrum/Intellectual Disability (ASID) database^14^) (Figure 2d).

While primary neural stem cells directly sourced from fetal telencephali represent the arguably most proximal model of human neurodevelopment, their availability is limited and hence they are ill suited for large-scale and iterative studies required to advance regulatory toxicology. We thus validated our transcriptomic findings on MIX N using two complementary neurodevelopmental models based on self-renewing sources of human induced pluripotent stem cells (iPSC): i) neuronal precursors differentiated from iPSC through short-term (7 days) Ngn2 over-expression^15^ and ii) apical progenitors (sourced at day 18 of culture) from iPSC-derived 3D cortical organoids that recapitulate human *in vivo* corticogenesis We confirmed in Ngn2-driven neuronal precursors the early activation of Nestin, PAX6 and SLC17A7 (VGLUT1) along with other defining neural markers (Extended data Figure 1b). Likewise, day 18 organoids upregulated neural progenitor genes (Extended data Figure 1b). Immunofluorescence confirmed expression of specific markers, including PAX6 and Nestin (Figure 2e), as well as, in organoids, the defining arrangement in ventricular zone-like structures lined by ZO-1 expressing cells (Figure 2f and Extended data Figure 1c).

Principal component analysis of the transcriptomes confirmed that organoids and Ngn2-driven precursors map closer to cortical than ganglionic eminence HFPNSC, confirming their dorsal fate (Extended data Figure 1d). In addition, organoids appeared more homogenous than Ngn2-driven lines. To assess the extent to which these systems recapitulate the effects seen in HFPNSC, we subjected them to the same exposures and profiled by RNA-seq the dose-response of the two clusters of DEGs identified above. Importantly, while we observed for both systems a highly significant overlap (p~3e-8 to 5e-12) with dysregulated genes in HFPNSC (Figure 2g), only organoids largely recapitulated the dose-response patterns seen for DEGs in HFPNSC (compare Figure 2i with 2b), thus representing a potentially transforming tool for large-scale regulatory toxicology.

Our validation of human iPSC-based neurodevelopmental models for assessing EDCs paves the way to the systematic evaluation of their toxicity across different human genetic backgrounds using representative iPSC collections from defined populations, for which we empirically derived the optimal transcriptome-based disease modeling designs^17^. Since MIX N was defined by verbal proficiency we thus reasoned that a genetic background conferring relative resilience of verbal skills would provide a rigorous proof of principle of the applicability of our findings across genetic backgrounds. To this end, we derived Ngn2-driven precursors and cortical organoids from iPSC lines harboring the 7q11.23 hemi-deletion that causes Williams-Beuren syndrome (WBS), a neurodevelopmental disorder characterized by cognitive weaknesses that selectively spare language abilities, and for which we previously uncovered disease-relevant dysregulation in iPSC and neural progenitors^18^. Using the same analytical approach outlined above (Figure 2j-k), we confirmed that even in this language-resilient genetic background, MIX N yields non-monotonic dose-responses with dysregulation of gene expression at low concentrations. In both experimental models, this dysregulation resulted in a significant enrichment of the DEGs for categories relevant for neurodevelopment, such as axonogenesis and synaptic signaling (Figure 2l-m). Interestingly, however, neither WBS organoids nor Ngn2-driven lines recapitulated the patterns of dose-response previously identified for the two clusters of DEGs from HFPNSC, which were recapitulated in control organoids, underscoring the effectiveness of iPSC-based models in uncovering genetic background-specific susceptibility to EDC exposure (Extended data Figure 1e).

Finally, we probed the cellular-level impact of MIX N on the SH-SY5Y human neuroblastoma line whose ability to undergo rapid differentiation *in vitro* (including formation of growth cones and elongated neurites with nodes) has established it as a relevant model in toxicology^19^. Image analysis of SH-SY5Y cultures treated with MIX N at 6 concentrations for 4d revealed a significant relative increase of more mature, extended node-bearing neurites (Figure 2n) and, consistently, increased expression of the growth cone-associated gene *GAP43* (Figure 2o), pointing to EDC-induced premature differentiation including dysregulated axonogenesis.

### MIX G disrupts human metabolic pathways

Following the same logic applied to MIX N, we evaluated the molecular impact of MIX G on two growth/metabolism-relevant human models, bone marrow-derived mesenchymal stem cells (adult MSCs) and iPSC-derived mesenchymal stem cells (iPSC-MSC) which display comparable molecular hallmarks (Figure 3a), allowing validation across experimental systems.

**Figure 3:**
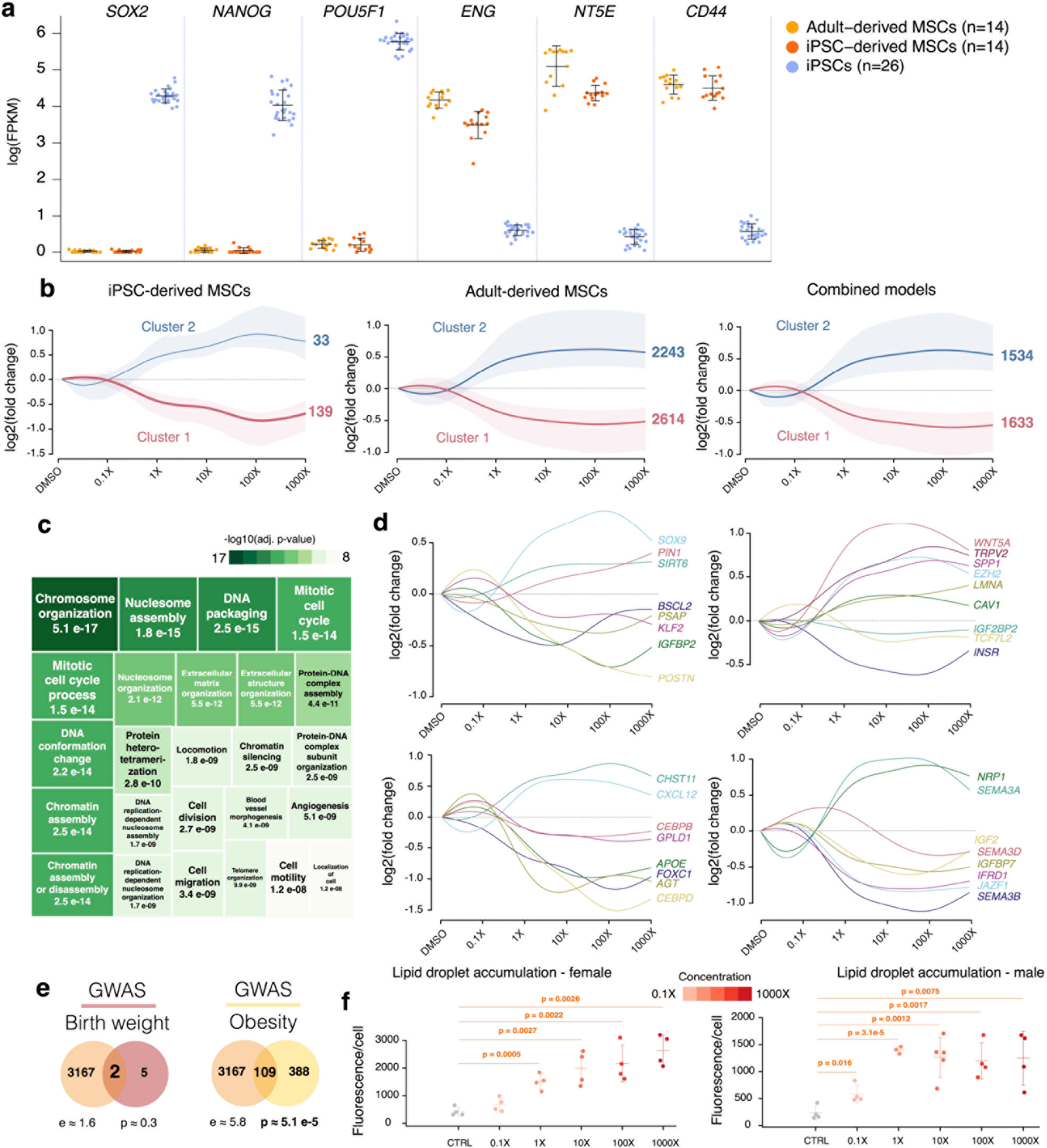
MIX G disrupts metabolic pathways in human MSCs. **a:** Log-transformed Fragments Per Kilobase of transcript per Million mapped reads (FPKM) of specific pluripotency and mesenchymal markers in MSC experimental systems and iPSC. **b:** Unbiased DEGs identified through categorical analysis for adult MSCs, iPSC-derived MSCs or both systems combined, clustered into major dose-responses patterns and plotted across the different MIX G dilutions (the size of the line is proportional to the number of DEGs in the cluster, shades represent 80% variance across samples). **c:** Treemap of the enriched Gene Ontology (GO) terms in the DEGs from the merged analysis of the two MSC systems. **d:** Dose-response patterns of DEGs that associated to adipogenesis and osteogenesis. **e:** Overlaps, enrichment and significance between unbiased mesenchymal DEGs and genes associated to alteration of birth weight and obesity. **f:** Quantification of lipid droplet accumulation in adult MSCs by Bodipy 493/503 staining upon treatment with the indicated concentrations of MIX G for 3 weeks. Values are normalised to nuclei count and representative of three independent experiments for each of the 2 donors, shown as mean and S.D. from 3 to 6 replicates of a single experiment.

Both MSC models were exposed to increasing concentrations of MIX G for 48h and profiled by RNA-Seq. In both systems DEGs grouped in two clusters, showing dose-dependent increases or decreases upon MIX G exposure (Figure 3b). Notably, clusters were very similar for the two different MSC sources but considerably different from those identified in the neurodevelopmental systems with MIX N, pointing to the robust capture of tissue-specific responses across experimental systems.

The unbiased DEGs from the merged MSC analysis were significantly enriched for GO terms related to chromatin regulation (Figure 3c), while DEGs in the two different clusters were enriched for extracellular matrix organization (downregulated genes) or cell cycle processes and chromatin regulation (upregulated genes) terms (Extended data Figure 1f). Importantly, EDC-dependent dysregulation included specific well-established genes linked to adipogenesis or osteogenesis (Figure 3d) and genes associated with birth weight and obesity through Genome Wide Association Studies (GWAS) (Figure 3e).

We probed a functional readout of these molecular alterations by exposing adult MSCs to the same doses of MIX G for 21 days and staining lipid droplets with Bodipy 493/503. In line with the epidemiological and transcriptional outcomes, MIX G significantly enhanced adipogenic differentiation at 1X concentration (Figure 3f) with increased PPARY expression, a master regulator of adipocytic differentiation, after 14 days of exposure (Extended data Figure 1g).

### Dissecting the impact of MIX N versus MIX G on human development

Despite having similar chemical compositions, MIX N and MIX G had been linked to different health outcomes on the basis of epidemiological evidence. We thus sought the molecular basis of this distinction by cross-exposure of representative models to the alternative mixture. HFPNSC were exposed to MIX G, using the same five dilutions as for MIX N. As expected, the two mixtures showed a general transcriptomic impact of similar magnitude (369 unbiased DEGs for MIX N versus 275 for MIX G) and with significant overlap of the DEGs (p~1e-3). Strikingly however, the mixtures showed marked differences in the affected genes, in particular with respect to ASD and ID associated targets that were only significantly enriched in MIX N DEGs (Figure 4a), consistent with the association of MIX N exposure to early verbal skills. Moreover, even those DEGs associated with ASD or ID that were altered by both MIX N and MIX G were impacted at the lowest doses of MIX N but only at the highest doses of MIX G (Figure 4b).

**Figure 4:**
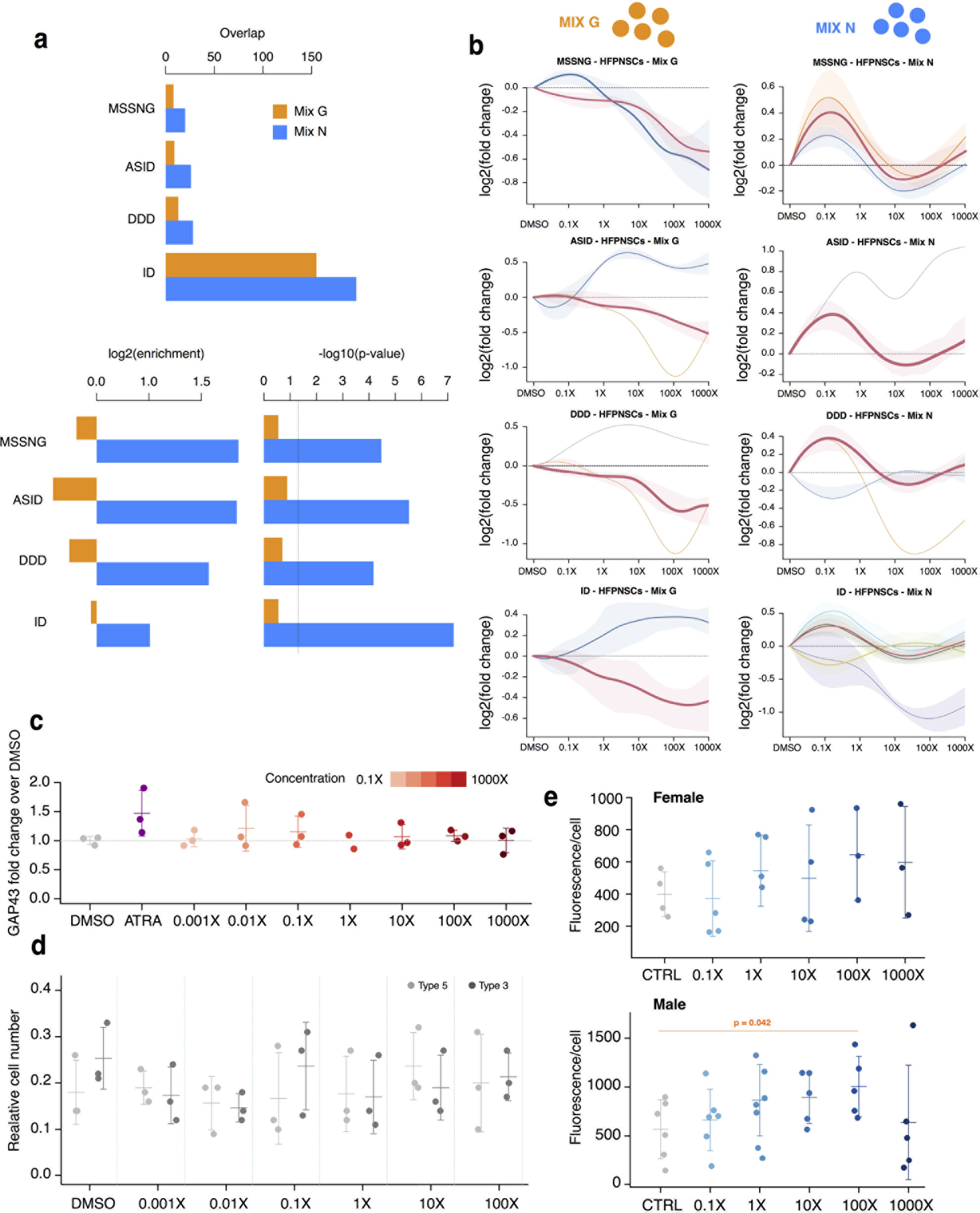
Differential impact of MIX N vs MIX G on human developmental systems. **a:** Overlaps, enrichment and significance between relevant gene sets and, respectively, MIX N- and MIX G-associated DEGs in HFPNSC. **b:** Comparison of the dose-response patterns, upon treatment with each mixture, for the union of DEGs, identified upon exposure to either of the two mixtures, that are associated to autism, intellectual disability or developmental disorders (the size of the line is proportional to the number of DEGs in the cluster, shades represent 75% variance across samples). **c:** Effect of MIX G on *GAP43* mRNA expression in SH-SY5Y cells. Treatment with 10 μM ATRA was included as a positive control. Data were quantified by the relative standard curve method and expressed as the relative amount of *GAP43* divided by the relative amount of *GAPDH*. Data are shown as mean α SD of 3 experimental replicates (except 1X G for which n=2). d: Relative number of cells with immature neurites (Type 3) or maturing neurites with nodes (Type 5) in SH-SY5Y cells treated with MIX G or vehicle (0.01 % DMSO) for 96 hours. Data are shown as mean ± SD of the relative amount of type 5 and 3 cells with three experimental replicates. **e:** For adult MSCs, lipid droplet accumulation was quantified using Bodipy 493/503 staining upon treatment with the indicated concentrations of MIX N for 3 weeks. Values are normalised to nuclei count and representative of three independent experiments for each of the 2 donors, shown as mean ± SD from 3 to 6 replicates of a single experiment.

Finally, to evaluate the specificity of MIX N and MIX G on cellular responses, we compared their effects on neurite outgrowth and *GAP43* expression in SH-SY5Y cells and on lipid droplet accumulation in adult MSCs, respectively. Contrary to MIX N (Figure 2n+o), MIX G had no effect either on cell morphology or *GAP43* expression (Figure 4c+d). Similarly, whereas adult MSCs exposed to MIX G showed significant increase in lipid accumulation already at 1X (Figure 3f), MIX N showed significant increase only at 100X and only in the male line (Figure 4e).

Together, these results provide experimental evidence of the mixture-to-phenotype dissection that had been originally only inferred at the population level, establishing the power of such integrated approaches for defining the molecular traces of EDC exposure across the population, organismal and cellular scales.

### MIX N and Mix G disrupt thyroid hormone signalling and related behaviour in developmental in vivo models

Having determined the markedly specific impact of MIX N and MIX G in the *in vitro* systems, we reasoned that *in vivo* key endocrine pathways might be disrupted by both mixtures but with specific downstream consequences. We thus probed the thyroid hormone (TH) axis as paradigmatic candidate for convergent dysregulation, given its essential roles in both brain development^20^ and metabolism^21^ and previous epidemiological^22,23^ and experimental^24^ evidence implicating the main chemical classes present in both mixtures as TH disruptors^25^.

The TH disrupting capacity of the mixtures was investigated using the *Xenopus* Embryonic Thyroid Assay (XETA), which serves as an endpoint assay for TH disruption at multiple levels (synthesis, transport and metabolism). TH signaling is well conserved across vertebrates; early stage Xenopus embryos are TH sensitive and metabolically competent^27^, making them a tractable model for assessing thyroid disruption. One-week old transgenic tadpoles, harboring a GFP thyroid responsive element construct, were exposed for 72h with or without TH. The mixture was renewed daily, and GFP expression levels quantified (Figure 5a+b). Interestingly, both mixtures altered TH availability in the brain, thereby providing insight into the potential adverse outcomes observed in human cohorts. Exposure to MIX N at concentrations 1X, 10X, 100X, and 1000X resulted in significant reduction in fluorescence at concentrations 1X and 10X. This decreased TH availability acquires human relevance when placed in the context of epidemiological results^28^, as slight changes in maternal TH levels during early pregnancy results in IQ loss and modified brain structure in offspring. Similarly, exposure to MIX G significantly decreased fluorescence at 10X and 100X (Figure 1c). To further investigate the effects of each mixture, neural gene expression of 3 day-exposed tadpoles was analysed by RT-qPCR. MIX N exposure decreased the expression of the TH receptor, *thra*, at 10X and 100X concentrations and the TH-transporter, *oatp1c1*, at 10X (Figure 5c). When tested with a 5 nM T3 spike, 1000X MIX N downregulated expression of 4 genes involved in the TH signalling pathway; *dio3*, *thibz*, *thrb* and *klf9* (Extended data Figure 3b). Exposure to MIX G also affected TH targets, significantly increasing *klf9* mRNA levels at 1X and 100X (Extended data Figure 2b), whilst significantly decreasing levels of *thibz* and *dio3*, at 1000X when tested with a5 nM T_3_ spike (Extended data Figure 3e).

**Figure 5:**
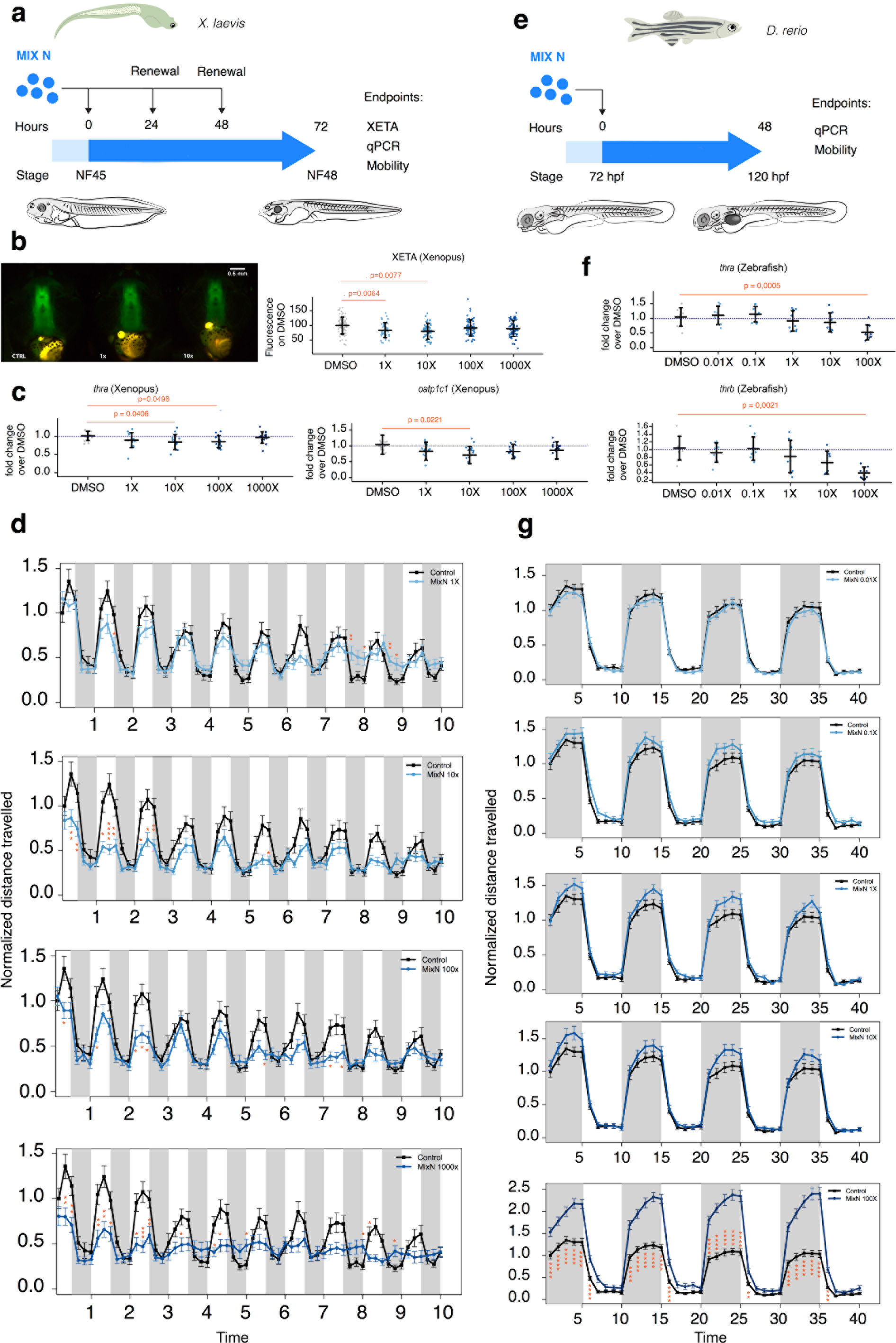
MIX N exposure disrupts thyroid hormone signaling and normal behavior both in *Xenopus laevis* and *Danio rerio*. **a:** The experimental setup for *X*. *laevis*. **b:** Thyroid disrupting effects were assessed using Xenopus Thyroid Embryonic Assay (XETA). Representative images of CTRL, 1X and 10X exposed tadpoles are shown on the left. Specific fluorescence in tadpoles’ head was quantified as relative fluorescence units normalized against CTRL (right). A pool of five independent experiments with >10 tadpoles per concentration per experiment is shown as mean ± SD. A significant decrease is found at 1X and 10X concentration. **c:** RT-qPCR following brain dissection for TH early-response target genes (*thra* and *oatp1c1*) in wild type tadpoles exposed to protocol (a). Shown is a pool of three independent experiments with mean ± SD (4 to 5 replicates per concentration per experiment). d: Mobility of exposed tadpoles was measured for 10 minutes in 30 sec light/30 sec dark cycles. Total distance travelled was analyzed over time. Shown is a pool of three independent experiments as mean ± SEM with 7-12 tadpoles per concentration per experiment. e: The experimental setup for *Danio rerio*. **f:** *D*. *rerio* embryos were exposed (protocol (e)), and RT-qPCR was performed on pooled whole larvae for TH early-response target genes: *thra* and *thrb*. Shown is a pool of the three independent experiments with mean ± SD (3 replicates per concentration per experiment). **g:** Mobility of exposed zebrafish embryos (120 hpf) was measured for 40 minutes in 5 min dark/5 min light cycles. Distance travelled was analyzed over time. Shown is a pool of the three independent experiments as mean ± SEM with 6 to 8 larvae per concentration per plate and 3 plates per experiment. Significance for both mobility experiments: parametric One-way ANOVA or nonparametric Kruskall-Wallis test. *P<0.05, **P<0.01, ***P<0.001, ****P<0.0001.

To confirm the impact of the mixtures on the thyroid system in an additional toxicologyrelevant *in vivo* model, we assessed expression of TH-related genes in zebrafish (*Daniorerio*) larvae. Zebrafish embryos were exposed to MIX N or MIX G for 48h (72 hpf-120 hpf) and expression levels determined by RT-qPCR (Figure 5e). MIX N exposure significantly decreased expression of both TH receptors, *thra* and *thrb*, at 100X (Figure 5f). Remarkably, MIX N exposure also downregulated the homologs of two particularly relevant DEGs affected in the human neurodevelopmental systems (Extended data Figure 3c) i) *kmt2d*, whose mutations cause Kabuki syndrome, a well-established neurodevelopmental disorder, encoding a histone H3 lysine 4 methyltransferase that competes with NCOR (a well-documented TH co-repressor) to regulate chromatin status and Notch targets^29^; and ii) *gatad2b*, encoding p66, a subunit of the NuRD chromatin remodelling complex whose interference relieves TH-mediated transcriptional repression^30^. MIX G exposure also affects TH signaling, significantly increasing expression of *thra* (Figure 1c), and deiodinases *dio1* and *dio2* as well as *thrb* at 0.1X and 1X (Extended data Figure 2e).

Finally, to establish the organismal-level impact of the mixtures on relevant behaviors, we used locomotor assays to asses TH-disrupting effects, given the prominent role of TH in maturation of the central and peripheral nervous system^20^. We analyzed light-induced locomotion in Xenopus tadpoles after 72h MIX N or MIX G exposure. Mobility was significantly decreased following exposure to MIX N at 10X, 100X and 1000X (Figure 5d) and MIX G at 1X, 10X and 100X (Extended data Figure 2c). In Zebrafish larvae, in which activity increases in darkness periods and abates in light^31^, behavioural effects were assessed after 48h exposure to MIX N and MIX G revealing significant induction of mobility during dark periods following exposure to 100X MIX N (Figure 5g). A tendency to hyperactivity was also seen at 10X, but this did not reach statistical significance. Similarly, mobility was significantly induced after 100X exposure to MIX G (Extended data Figure 2f). The contrasting responses to mixture exposure between the two models is expected to be related to documented differential timings of TH-dependent processes within the maturing brain and periphery for the two species^32^.

Taken together, these results demonstrate TH-disrupting effects of MIX N and MIX G at molecular and physiological levels, at concentrations measured and associated with adverse outcomes in humans.

### The thyroid axis is a key site of vulnerability to MIX N and MIX G

Key genes involved in TH signaling, such as *thra*, *thrb* and *klf9*, were dysregulated in both *Danio rerio* and *Xenopus laevis* embryos. We therefore selected these as starting points to visualize pathways linking TH-signaling to the DEGs identified in HPFNSC exposed to MIX N and G (Figure 6), using the Genomatix Pathway System (GEPS) program. This software determines interactions through both publically available and inhouse data^33^, ^34^ Of the 1,848 DEGs identified in HPFNSC under MIX N exposure, we focused on the 185 genes implicated in neurodevelopmental disorders. THRA, THRB or KLF9 (Figure 6, green box) were directly of indirectly linked to 12 of these DEGS (Figure 6, blue box) (see Supplementary Table 1 for interactions), including genes encoding three main THR-comodulators, namely, *NCOA1*^35^, *CREBBP*^36^ and *SIN3A*^37^ which are associated with neurodevelopmental delay when mutated. These results highlight specific genes as critical hubs of disruption by common environmental chemicals that are conserved across toxicological models^24,38^. TH signaling is also an essential player in growth^39^ and metabolic control^21^. Both GH and IGF pathways interact with and are controlled by TH at multiple levels^40^. We therefore applied a similar approach to the DEGs identified in adult MSCs following exposure to MIX G. Twenty one key DEGS were identified as regulated by either THRA, THRB or KLF9 (Figure 6, orange box). For instance, THRB binds TP53^41^, a transcription factor that directly inhibits ERCC6 activity^42^ and mutations of ERCC6 are associated with slow growth^43^. THRB also binds SRC^44^, which directly phosphorylates CAV1^45^. Mutations in CAV1 correlate with insulin resistance^46^. Further, KLF9 is a direct regulator of TCF7L2^47^, a diabetes-susceptibility gene^48^.

**Figure 6:**
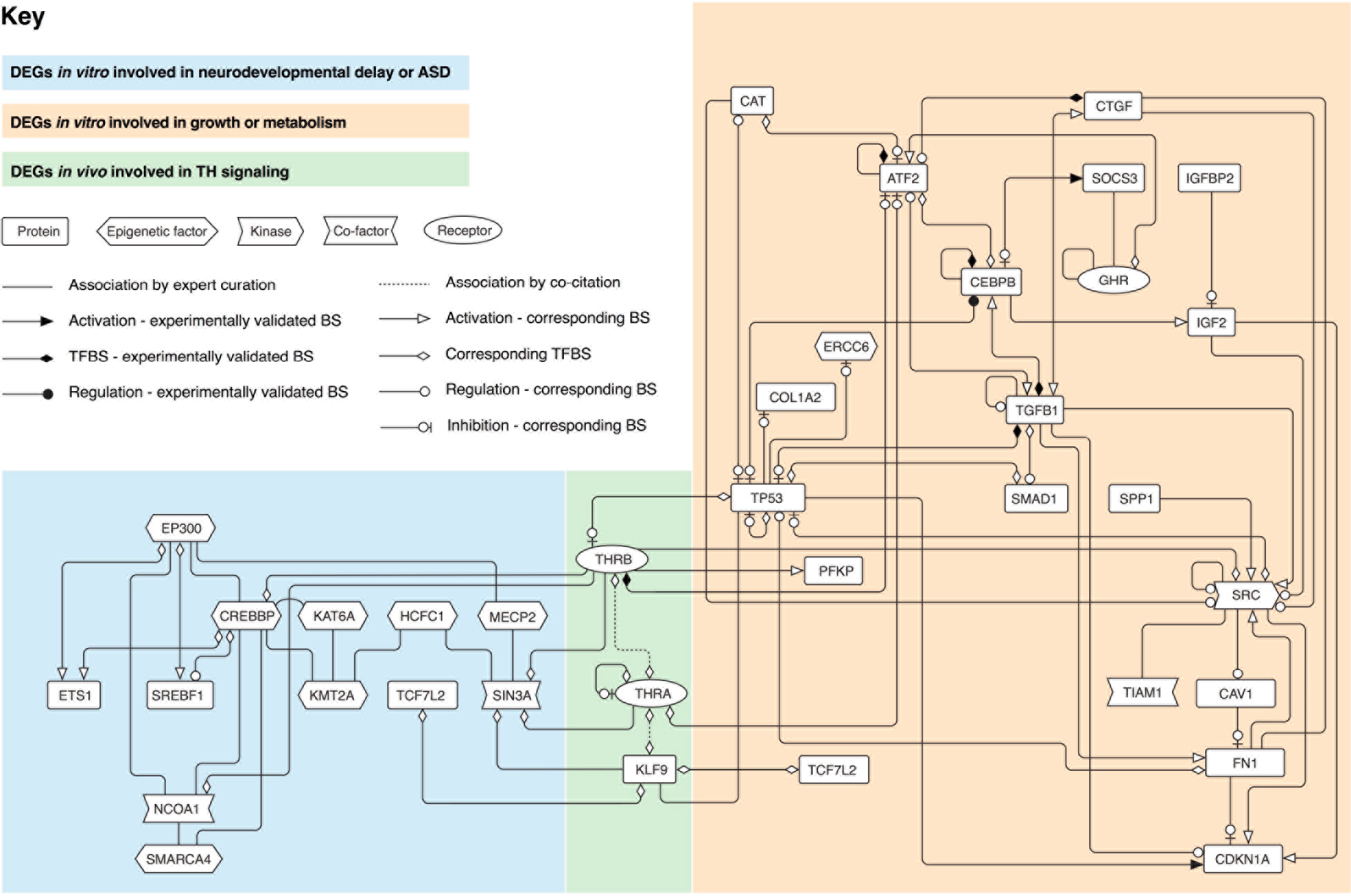
Thyroid hormone receptor (THRA and THRB)- and KLF9-linked pathways regulate Differentially Expressed Genes (DEGs) identified in human models. Regulatory pathways were generated by the Genomatix GEPS program that connects differentially expressed genes. Green box: starting from DEGs identified *in vivo* in *Xenopus* and *Danio rerio*, interactions were established with disease-linked DEGs identified in HFPNSC. Orange box: DEGs identified in adult MSC and iPSC-derived MSC human lines related to growth or metabolism. Blue box: DEGs identified in HFPNSC and implicated in neurodevelopment or ASD. TFBS stands for transcription factor binding site and BS for binding site.

## Discussion

The current vision for improving regulatory decision making relies on the transforming potential of high throughput and high content data to elucidate and quantify the molecular, cellular and organismal responses to chemicals^49^. In the context of chemical regulation most authorities, including the Organisation for Economic Co-operation and Development (OECD), recommend integrated approaches to testing and assessment (IATA) that incorporate results from multiple methodologies. Emphasis is placed on molecular initiating events (MIE) that lead to physiologically measurable adverse outcome pathways (AOP). Here we first identify the adverse outcomes (language delay or low birth weight) in humans, then proceed to determine the prenatal chemical mixtures associated with these outcomes in children and, finally, establish the causative molecular and cellular impacts using *in vitro* and *in vivo* models. By making EDC mixtures experimentally tractable as the ‘real life’-relevant unit of exposure, these complementary methodologies allowed us to uncover the gene networks specifically altered by neurodevelopment- or growth-targeting EDC mixtures and define thyroid function as a key and unifying axis of vulnerability to both mixtures. Furthermore, by establishing the value of human reprogrammed models based on self-renewing sources, we both expand their reach to regulatory toxicology and enrich the latter experimental human insight.

Together, this approach allowed us to define dysregulated gene networks, identified and validated through complementary methods, that can be exploited to link, mechanistically, MIEs to AOPs. As such we expect the methodology and approach to be broadly applicable within the new regulatory frameworks globally.

## METHODS

Detailed protocols and methods descriptions including references can be found in the supplementary information (SI).

### Exposure assessment

Using data from the Swedish Environmental Longitudinal Mother and child Asthma and allergy (SELMA) pregnancy cohort^8^ (described in SI), mixtures of prenatal EDC exposures of relevance for health outcomes in children were identified. Exposure was measured in urine and serum taken in week 3-27 of pregnancy (median week 10, and 96% of the samples were taken before week 13). First morning void urine samples were analyzed for 10 phthalate metabolites (Mono-ethyl phthalate (MEP), metabolite of DEP; Mono-n-butyl phthalate (MnBP), metabolite of DBP; Monobenzyl phthalate (MBzP), metabolite of BBzP; Mono-(2-ethylhexyl) phthalate (MEHP), Mono-(2-ethyl-5-hydroxylhexyl) phthalate (MEHHP), Mono-(2-ethyl-5-oxohexyl) phthalate (MEOHP), Mono-(2-ethyl-5-carboxypentyl) phthalate (MECPP), metabolites of DEHP; Mono-hydroxy-iso-nonyl phthalate (MHiNP), Mono-, oxo-iso-nonyl phthalate (MOiNP), Mono-carboxy-iso-octyl phthalate (MCiOP), metabolites of DiNP); and alkyl phenols including Bisphenol A (BPA) and Triclosan (TCS)). Serum samples were analyzed for 8 perfluorinated alkyl acids (perfluoroheptanoic acid (PFHpA), perfluorooctanoic acid (PFOA), perfluorononanoic acid (PFNA), perfluorodecanoic acid (PFDA), perfluoroundecanoic acid (PFUnDA), perfluorododecanoic acid (PFDoDA), perfluorohexane sulfonate (PFHxS) and perfluorooctane sulfonate (PFOS)) as described in SI and publications therein.

### Health examinations

For a measure of metabolism and growth in the children, we used birth weight data from the Swedish national birth register. For a measure of neurodevelopment we used data from a routinely made language screening of the children when they were 30 months old. Language development was assessed by nurse’s evaluation and parental questionnaire, including the number of words the child used (<25, 25-50 and >50). A main study outcome was parental report of use of fewer than 50 words, termed language delay (LD) corresponding to a prevalence of 10%.

### Biostatistic analyses

Weighted quantile sum (WQS) regression^9^, adjusted for covariates, was used to establish associations between mixture exposures and lower birth weight or language delay in children (see SI). In short, WQS regression is a strategy for estimating empirical weights for a weighted sum of concentrations most associated with the health outcome. The results are a beta coefficient associated with the weighted sum (estimate, SE and p value) and the empirical weights (which are constrained to sum to 1). The components most associated with the health outcomes have non-negligible weights, and were treated as “bad actors”.

Next, we estimated the equivalent daily intake (DI) of “bad actors” measured in the urine (i.e., phthalates and alkyl phenols), and estimated serum concentrations from the DI for these urinary measurement-based compounds (see SI). Finally, we used the geometric means, on a molar basis, for either the measured (PFAAs) or estimated serum levels (phthalates and alkyl phenols) and established mixing proportions to prepare one mixture associated to low birth weight (MIX G) and one associated to language delay (MIX N). These two mixtures were used in the experimental validation.

### Composition of the mixtures

The chemicals needed for the mixtures were obtained from commercial custom synthesis laboratories or vendors BPA, Dimethylsulfoxide (DMSO), MBzP, PFHxS, PFNA, PFOA and PFOS were obtained from Sigma-Aldrich Inc. (St. Louis, MO, USA). Triclosan was purchased from Dr. Ehrenstorfer (Augsburg), MEP and MiNP were obtained from Toronto Research Chemicals (North York, ON, Canada). MBP and MEHP were purchased from TCI, Tokyo Chemical Industry Co., Ltd (Japan). For MIX N, 1M solutions in DMSO were prepared using, MEP, MBP, MBzP, MiNP, BPA, PFHxS, PFNA, PFOS. For MIX G, 1M solutions in DMSO were prepared using, MEP, MBP, MBzP, MEHP, MINP, Triclosan, PFHxS, PFOA, and PFOS. Thereafter, the 1M solutions were mixed in proportions as described SI.

### Cell lines

Human induced pluripotent stem cells have been previously validated in Dr. Giuseppe Testa's laboratory. hiPSCs were cultured on plates coated with human qualified Matrigel (BD Biosciences) diluted 1:40 in DMEM-F12, in the following cell culture media: mTeSR1 medium (StemCell Technologies). Two iPSC lines, CTL1R-3 and WBS2-C2 were used for the experiments.

Human foetal primary neural stem cells (HFPNSC) were provided by Dr. Steve Pollard’s laboratory. They were derived from the cortex of post-conception week 11, male embryo and from the ganglionic eminence of post-conception week 8, male embryo.

The SH-SY5Y human neuroblastoma cell line was provided by Dr. June L. Biedler and cultured as described in SI.

Adult human bone marrow derived mesenchymal stem cells (hMSCs) from 2 donors were a kind gift of Dr. Katarina Leblanc (Center of Hematology and Regenerative Medicine, Department of Medicine, Karolinska Institutet, Sweden) and were cultured in Dulbecco’s Modified Eagle’s medium (DMEM, Gibco® by Life technologies) supplemented with 10% heat-inactivated fetal bovine serum (FBS) (Gibco® by Thermo Fisher Scientific) (see also SI). Human indcuded pluripotent stem cell-derived mesenchymal stem cells (hiPSC-MSCs) were obtained and cultured from neural crest stem cells derived from iPSCs in Dr. Giuseppe Testa’s laboratory (IEO, Milan Italy) as described in SI and references therein.

### Cell differentiation and treatment

For the induction of neuronal differentiation of iPSC, two protocols were followed. For generating Ngn2-driven neuronal precursors the protocol described by Zhang et al.,^16^ was followed and some improvements were made in Dr. Giuseppe Testa’s laboratory (IEO, Milan Italy). Clonal cell lines were derived from the infection by splitting the infected bulk and plating single cells in separate wells of a 96-well plate. This was achieved by sorting DAPI-negative cells in the plate and waiting for them to grow as colonies, then passaging them progressively increasing the dish size, until they were safely frozen. The differentiation protocol was carried out without using astrocytes inside the culture plate giving us the opportunity to avoid the important loss of sequencing reads that usually account for astrocytic cells. To overcome their absence, culture media was conditioned for 48 hours in mouse astrocytes coated dishes before using it for differentiating cells. In transcriptomic experiments the stable lines were seeded in 6 well plates (2×10^5^ cells/well) and then differentiated toward the neuronal lineage following the published protocol steps. At the 5^th^ day of differentiation, DMSO or MIX N in 5 different concentrations were diluted in the culture media and added to the cells for 48 hours.

For generating human cortical organoids the protocol described by Pasca et al.,^17^ was followed and some improvements were made in Dr. Giuseppe Testa’s laboratory (IEO, Milan Italy) to improve the apoptotic core that usually characterize the inner part of the organoids. The GSK-3b inhibitor CHIR99021 was added to the culture media from the time of generation of embryoid bodies. Since the 14^th^ day of differentiation, dishes were positioned on a shaker inside the incubator. For transcriptomic experiments, on the 16^th^ day of differentiation 3 organoids were plated in each well of a 6 well ultra-low attachment plate, DMSO or MIX N in 5 different concentrations was diluted in the culture media and added to the organoids for 48 hours.

HFPNSC were seeded in 6 well plates. When confluency was reached DMSO or MIX N or MIX G in 5 different concentrations was added to the culture media and used to culture cells for 48 hours.

For experiments with SH-SY5Y, the cells were seeded at a density of 3600 cells/cm^2^ in complete culture medium. MIX G and N were added at the time of seeding (0 h) and 48 h later before harvest at 96 h. Ten μM all-trans retinoic acid (ATRA) was added 24 h after seeding as a positive control.

For the induction of adipogenic differentiation, cells were seeded in 96 well plates (2×10^4^ cells/cm^2^) and allowed to expand until they reached 80% confluence. Two days before MIX G/N treatment or initiation of adipogenic induction, growth media was replaced by treatment media consisting of DMEM supplemented with 10% charcoal stripped FBS, 1% penicillin/streptomycin and 2% glutamine. Subsequently, treatment was performed for 21 days whereby medium was changed twice a week.

### RNA sequencing

Total RNA was isolated with the RNeasy Micro Kit (Qiagen, Hilden, Germany) according to the manufacturer’s instructions. RNA was quantified with Nanodrop and then the integrity was evaluated with Agilent 2100 Bioanalyzer (only if the quality ratios were not optimal after Nanodrop analysis). TruSeq Stranded Total RNA LT Sample Prep Kit, Illumina was used to run the library for each sample. Spike-ins (ERCC RNA Spike-in control Mixes, Life Technologies) were added to the sample before proceeding with the protocol to validate the process. Sequencing is performed with the Illumina HiSeq 2000 platform, sequencing on average 10 millions 50bp paired-end reads per sample.

### RNA-seq data analysis

RNA-seq quantification was performed directly from the reads using Salmon 6.1, using the hg38 Refseq annotation complemented with the sequences of ERCC spike-ins. Only genes with at least 20 reads in each of at least 2 concentrations of the mixture using the same cell lines were included for further analysis; small (<200nt) genes, ribosomal RNA genes, and fusion genes were excluded. Differential expression analysis was performed on the estimated counts after TMM normalization with edgeR v.3.12.1 using a likelihood ratio test on the coefficients of a negative binomial model including the genetic background and the mix’ concentration (i.e., ~line+concentration). For the first, unbiased analysis, the concentration was treated as a categorical variable (i.e., converted to factor), and tested for any non-zero coefficient. Genes identified through this method were then kmeans-clustered on the basis of their smoothed fold change upon each concentration (using the NbClust R package to determine the consensus number of clusters). The mean smoothed fold change pattern for the main cluster(s) were then used as an independent continuous variable for a second test retrieving more genes following the same pattern.

### Enrichment analysis

Gene Ontology (GO) enrichment analyses were performed with the goseq R package, including correction for eventual RNA-seq transcript-length bias and excluding genes without annotation. Terms with at least 10 but no more than 1,000 associated genes were considered, and Fisher’s exact test was used. The tested genes (excluding small and lowly-expressed genes) were used as a background. Parent terms with significantly enriched children terms were filtered out to improve the specificity of the enrichments. Unless stated otherwise, other enrichment tests were performed using the hypergeometric test.

### Immunohistochemistry for neuronal systems

Protocols and antibodies used for immunohistochemistry are described in SI.

### RNA extraction and quantitative PCR

Protocols and primer sequences uses for RNA extraction and qPCR analyses for all systems are described in SI.

### Neurite morphology in SH-SY5Y cells

Cells were seeded in 35 mm dishes and exposed to 0.01 % DMSO or the indicated concentrations of MIX N or MIX G, or ATRA as described above. After 96 hours the living non-fixated cells were examined under an inverted phase contrast microscope (Leica DMI6000B, Germany) and ten to fourteen fields per condition were photographed. Morphological assessments of 150 to 200 cells per condition and experiment using the ImageJ software (NIH shareware, v 1.49) were made with the observer blinded to treatment. The cell morphology was judged according to criteria scoring for sprouts, neurites (protrusions longer than one cell body diameter) without nodes and more mature neurites containing nodes. Statistical analysis for dose-response patterns using ANOVA was done for individual neurite types and ratios thereof.

### Bodipy 493/503 and Hoechst 33342 staining

Cells were seeded in black-walled 96 well plates with μCLEAR bottom (Greiner Bio One) and exposed to DMSO or the indicated concentrations of MIX G or N as described above. Staining was performed as described in SI and references therein. Images were acquired immediately using the Image Xpress Micro High-Content Analysis System (Molecular Devices, Sunnyvale California USA). Images were taken in FITC and DAPI channel at 10x magnification, at 16 sites per well. Images were further analyzed with the MetaXpress High-Content Image Acquisition and Analysis software (Molecular Devices, Sunnyvale California USA). Using the Transfluor HT analysis module nuclei were counted and lipid droplets were quantified. Integrated granule intensity per image was normalized to nuclei count. Mean values of all images from six replicate wells were compared among different treatments.

### X. laevis rearing and strains

The used methods, care and treatment of *Xenopus laevis* in this study was in accordance with institutional and European guidelines (2010/63/UE Directive 2010) and the local ethic committee (Cometh: Comité d—Ethique en matière d’expérimentation animale) under the project authorizations No. 68-039. Heterozygous *X*. *laevis* tadpoles used for the XETA (Xenopus Embryonic Thyroid Assay) were obtained by crossing adult homozygous *Tg(thibz:eGFP)* with wild type animals. Wild type (naive) tadpoles were obtained by crossing wild type (WT) adults. Sexually mature males were mated with females that were injected the day before with 500-800U of human chorionic gonadotropin (Chorulon, France). Selected tadpoles were sorted according to the developmental stages.

### Exposure of X.laevis

Triiodothyronine (T3, Sigma-Aldrich,Saint-Quentin Fallavier, France) was prepared in 70% milliQ Water, 30% NaOH, at 10^−2^M, aliquoted in volumes of 100μL in 1.5mL low binding Eppendorf (100% polypropylene) tubes and stored at -20°C until use. For XETA and qPCR analyses, 15 tadpoles per well (stage NF45 e.g., 1 week old) were incubated in a 6-well plate (TPP, Switzerland). Each well contained 8 mL of exposure solution made of Evian water (or T3 5.10-9M prepared in Evian) and DMSO (containing or not mixtures at different concentrations). These exposure solutions were extemporaneously prepared in Greiner (France) polypropylene tubes (50 mL tubes) and transferred in the wells Final DMSO concentration was 0.01% in all treatment groups. The exposure time was 72 hours in the dark at 23μC with 24-hour renewal. MIX N and MIX G screenings were done in presence or absence of T3 5.10-9M except for mobility experiments (only absence of T3). After 72h exposure, tadpoles were rinsed and tested for mobility or anesthetized with 0.01% tricaïne methanesulfonate (MS222, Sigma-Aldrich, Saint-Quentin Fallavier, France) either for fluorescent screening (XETA) or euthanized in 0.1% MS-222 for brain gene expression analysis.

### Image Analysis for Xenopus Embryonic Thyroid Assay (XETA)

Images were captured for fluorescence quantification. All pictures of a group were stacked, and processed as described^24^. Five independent experiments were performed for each mixture providing comparable results. GraphPad Prism 7 software was used for statistical analysis. All values were normalized (100%) to either CTRL group or T3 when mixtures were tested in co-exposure. Results are expressed as scatter dot blots with mean +/- SEM. A d’Agostino and Pearson normality test was carried out to determine distribution of values in each of the exposure groups. If normal distribution was found, a one-way ANOVA and Dunnett’ post-test was applied. If one of the compared groups did not pass the normality test, a Kruskal-Wallis test with Dunn’s post-test was applied. All groups were compared to the appropriate control group.

### X.laevis mobility

Wild type *X.laevis* embryos NF45 underwent 72h mixture exposure as described above. Mobility was recorded by the DanioVision (Noldus, Wageningen, The Netherlands) behaviour analysis system. Tadpoles were first rinsed and placed individually in 12-well plates (TPP, Switzerland) filled with 4 mL of Evian and put under the infrared camera. Tadpoles had 5 minutes in dark to accommodate before starting the protocol. Light inside the box was turned -on and -off at a regular 30 second interval, giving the tadpoles an external stimulus. The total distance travelled by each tadpole within the well was recorded for a total of 10 minutes. Analysis was done with EthoVision software (11.5, Noldus, Wageningen, The Netherlands). Normalization was done for each experiment on the CTRL value of distance done after 10 first sec. Three independent experiments were done with 7<n<12 per condition per experiment. A pool of the three experiments is presented. Differences between CTRL and different mixture concentrations were analysed using non-parametric Kruskal Wallis’ test followed by Dunn’s post test for each time point or with parametric one-way ANOVA with Dunetts post test. Differences were considered significant at p < 0.05 (*), p<0.01 (**), p<0.001(***) and p<0.0001 (****).

### Zebrafish husbandry

All fish were treated in accordance with Swedish ethical guidelines with the ethical permit (Dnr 5.2.18-4777/16) granted by the Swedish Board of Agriculture. AB-strain zebrafish (*Danio rerio*), obtained from SciLifeLab (Uppsala, Sweden) were kept in a recirculating ZebTEC system (Tecniplast, Italy) at the University of Gothenburg. Fish were maintained at 26 °C with a 14:10 h light/dark cycle and fed *ad libitum* two times daily. Before embryo collection, two adult males and two females were placed in breeding tanks, separated by a transparent barrier and left overnight. The barrier was removed shortly before onset of light the next morning and fish were allowed to breed. Fertilized eggs were collected within 60 min of spawning, rinsed and kept at 28°C in autoclaved zebrafish embryo medium with daily medium changes until exposure was initiated. Embryo medium consisted of 245 mg/L MgSO_4_.7H_2_O, 20.5 mg/L KH_2_PO_4_, 6 mg/L Na_2_HPO_4_, 145 mg/L CaCl_2_·2H_2_O, 37.5mg/L KCl and 875 mg/L NaCl in milliQ Water.

### Exposure of zebrafish

Healthy embryos from a minimum of three different breeding pairs were selected and randomly mixed for exposures to account for inter-population variability (OECD guidelines 236, 2013). The MIX N and G exposure solutions were prepared by serial dilution in glass Erlenmeyer flasks and the final dimethylsulfoxide (DMSO) concentration in all treatments was 0.01%. At 72 hpf, embryos were moved to glass petri dishes containing 30 mL exposure solution (MIX G/N or vehicle control in embryo medium) and incubated for 48h at 28°C with a 14:10 h light/dark cycle. Three replicates with 20-25 embryos per concentration were used and each experiment was independently repeated 3 times. After exposure, embryos were moved to 48-well plates and tested for mobility or collected for gene expression analysis.

### Zebrafish larvae mobility

Mobility of 5 dpf larval zebrafish after 48h exposure (72 hpf-120 hpf, as described above) was recorded with the View Point® automatic behaviour tracking system (ViewPoint Life Science, Montreal, CN) and an infrared camera. Larvae were transferred to individual wells in 48-well plates with 500 μl solution per well. Each plate contained individuals from each exposure concentration and was considered to be a technical replicate. After 15 minutes of initial acclimatization in light, locomotion was induced with alternating light/dark cycles (5 min/5 min)^26^ Distance travelled was recorded for a total of 40 minutes and analysed with the View Point® Zebralab software (ViewPoint Life Science, Montreal, CN). Values were normalized on the Control group value of distance travelled during the first 60 sec. For every exposure concentration, three independent experiments with three technical replicates (6 to 8n per replicate) each were performed. A pool of the three experiments is presented. Differences between Control and mixture concentrations were analysed with nonparametric Kruskal Wallis’ test or one-way ANOVA for each time point. Differences were considered significant at p < 0.05 (*), p<0.01 (**) and p<0.001(***).

### Gene Network Analyses

185 from a total of 1848 identified DEGs in HFPNSC were manually selected through literature search to be implicated in neurodevelopmental delay or ASD. 111 from a total of 3617 identified DEGs in both adult MSCs and iPSC-derived MSCs were selected to be important in growth or metabolism. *Thra, thrb and klf9* were used as starting points to possibly connect the above described DEGS from different *in vitro* systems in their respective fields. For connections analyses, the Genomatix software was used (http://www.genomatix.de) combining mining sources such as MatInspector and Genomatix Pathway Systems (GePS). In GePS, genes were mapped into networks based on the information extracted from public databases. For the analyses ‘expert level’ was used to generate the network. Here gene pairs are noted if they are manually curated interactions from Genomatix experts reviewing the original literature. The MatInspector software utilizes a library of matrix descriptions for transcription factor binding sites to locate matches in DNA sequences. For interactions see Supplementary Table 1.

**Supplementary Information** is linked to the online version of the paper

## Acknowledgments

The authors wish to dedicate this work to the memory of Prof. Bo A.G. Jönsson (1960 – 2016), whose dedication and knowledge was instrumental to this project. This work received funding from the European Union’s Horizon 2020 research and innovation programme under grant agreement No 634880, EDC-MixRisk (AB, JR, CGB, EN, ⍰BJ, CR, MJ, JBF, ML, BD, WK, JS, LB, EK, HK, GT), the European Research Council (ERC) (DISEASEAVATARS 616441 to GT); the EPIGEN Flagship Project of the Italian National Research Council (CNR)(GT); the ERANET-Neuron grant from the Italian Ministry of Health (AUTSYN)(PLG); the Umberto Veronesi Foundation (PLG); the Italian Ministry of Health (Ricerca Corrente Grant to GT); the Telethon Foundation (Grant GGP13231B to GT); Science Without Borders Program (CAPES, Brazil) (RBB); the Swedish Research Council Formas (GB, JR); Centre National de la Recherche Scientifique (CNRS) (JBF, BD), Muséum National d’Histoire Naturelle (MNHN)(BD, CNRS).

### Author Contributions

C.G.B is principal investigator for the SELMA study and responsible for the biostatistical modelling together with C.G. N.C. carried out all the experiments with human neurodevelopmental systems, including neuronal differentiation, EDC exposure, RNA-Seq libraries; P.-L.G designed the analytical strategy of transcriptomic data; G.D. designed and edited the figures; G.D and M.Z. generated Ngn2 monoclonal lines; N.C. and G.B. carried out exposure experiments with iPSC-MSC and adult MSC and performed RNA-seq; G.B. set up and performed adipocyte differentiation, lipid droplet assessment and RT-qPCR in adult MSC; V.M.K. established the imaging platform for lipid droplet accumulation quantification; F.R. planned, carried out and analysed the experiments with the SH-SY5Y neuroblast model including EDC exposure, analyses of morphology and gene expression; B.S. contributed to the planning and data analyses for the SH-SY5Y neuroblast model. E.N. planned, supervised the work and analysed the data in the SH-SY5Y model. P.-L.G and N.C. performed RNA-Seq bioinformatic analysis on both neurodevelopmental and mesenchymal systems; A.L.T., S.T. and N.C. set up the cortical organoids protocol; A.L.T. performed immunofluorescence of Ngn2-driven neuronal precursors and cortical organoids; R.B.B. and S.P. provided human foetal primary stem cells and performed immunofluorescence for them; F.C. and M.La. set up astrocyte-free Ngn2 differentiation protocol; L.B. carried out all the experiments with *Danio rerio*, including EDC exposure, RT-qPCR and mobility assay; L.B and J.S analysed the *Danio rerio* data; M.Le. carried out all the experiments with *Xenopus laevis*, including EDC exposure, XETA, RT-qPCR and mobility assay; B.D, J.B.F, M.Le. analysed the *Xenopus* data; M.Le. made the genomatix figure; N.E.C. and G.T. attended to the bioethical issues of the project; A.B., P.D., M.J., W.K, E.K., J.B.F, H.K., M.O., P.R., C.R. and O.S. contributed to the study design, discussions and critical reading of the manuscript; G.B., C.G.B., N.C., B.D., P.L.G., M.Le., L.B., C.J., J.R. and G.T. wrote the paper; C.G.B., B.D., J.B.F., C.G., E.N., J.R., J.S. and G.T. conceived, designed and supervised the study.

### Author Information

The authors declare no competing financial interests.

The datasets generated during and/or analysed during the current study will be available in the GEO repository.

